# Improving animal behaviors through a neural interface with deep reinforcement learning

**DOI:** 10.1101/2022.09.19.508590

**Authors:** Chenguang Li, Gabriel Kreiman, Sharad Ramanathan

## Abstract

Artificial neural networks have performed remarkable feats in various domains but lack the flexibility and generalization power of biological neural networks. Given their different capabilities, it would be advantageous to build systems where both network types can synergistically interact. As proof-of-principle, we show how to create such a hybrid system and harness it to improve animal performance on biologically relevant tasks. Using optogenetics, we interfaced the nervous system of the nematode *Caenorhabditis elegans* with a deep reinforcement learning agent, enabling the animal to navigate to targets and enhancing its food search ability. Agents adapted to strikingly different sites of neural integration and learned site-specific activations to improve performance on a target-finding task. The animal plus agent displayed cooperative computation and generalized to novel environments. This work constitutes a demonstration of how to improve task performance in animals using artificial intelligence interfaced with a nervous system.

---

Artificial and biological neural networks differ in fundamental ways. Artificial neural networks can be trained to fit complicated functions using human-specified scoring metrics and have been used to accomplish a broad array of computational tasks^1^. However, artificial intelligence algorithms often fail to generalize, and may not perform well when applied to problems that are even slightly different from the ones on which they were trained^2^. Biological neural networks, on the other hand, have evolved to perform computations that help animals generalize to new and changing environments. The complementary strengths of artificial and biological neural networks raise the question of whether they can be integrated into a system that can not only compute information in a directed way but can also improve behavior while generalizing to novel situations.

Previous works have attempted to use direct neural stimulation to improve performance on a variety of tasks, relying on manual specification for stimulation frequencies, locations, dynamics, and patterns^3–11^. A central difficulty with these approaches is that manual tuning is highly impractical, as activation patterns for a given task and set of neurons are often unknown^6^, and there is a combinatorial explosion of stimulation parameters to test. In addition, effective patterns can vary depending on which neurons are targeted and on the animal itself^12, 13^. Thus, even though technologies for precise neuronal modulation exist^14, 15^, there still lies the challenge of how an artificial intelligence algorithm can systematically and automatically learn strategies to activate a set of neurons to improve a particular behavior^16–20^.

Here we addressed this challenge using deep reinforcement learning (RL), assessing whether RL can autonomously integrate with an animal’s nervous system to improve behavior. In an RL setting, an agent collects rewards through interactions with its environment. By leveraging deep neural networks, RL algorithms have been able to successfully discover complex sequences of actions to solve a wide set of tasks^21–31^. These past successes relied on reward signals to train algorithms, a framework that can be readily adapted to biologically relevant goals, such as finding food or mates. Consequently, an RL-based approach has the potential to handle the main computational problems in behavior improvement through neuronal stimulation.

We interfaced an RL agent with the nervous system of the nematode *C. elegans* using optogenetic tools^14, 17^. In a natural setting, *C. elegans* must navigate variable environments to avoid danger or find targets like food. Therefore, we aimed to build an RL agent that could learn how to interface with neurons to assist *C. elegans* in target-finding and food search. We tested the agent by connecting it to different sets of neurons with distinct roles in behavior. The agents could not only successfully couple with different sets of neurons to perform a target-finding task, but could also generalize the task to improve food search across novel environments in a zero-shot fashion, that is, without any prior training. This ability to generalize performance to novel environments is an important feature in natural behaviors and was achieved by augmenting the animal’s native nervous system with artificial neural networks.

## Connecting the nervous system to AI and training agents

We used a closed-loop setup to couple an RL agent to an animal’s nervous system (Figure 1A, B). We first formulated target-finding as an RL problem by defining a dense reward that increased with an animal’s proximity to a target (Figure 1C; Methods). The RL agent’s environment consisted of a ∼1 mm adult animal and a 4 cm-diameter arena on an agar plate. Observations of the environment were given to the agent through a camera at 3 Hz. Features were automatically extracted from each camera frame to track the animal’s center of mass (*x*_*t*_, *y*_*t*_) and its head and body angles 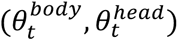 relative to the +*x*-axis. We took polar coordinates of the angle measurements so that for every frame at time *t*, we defined an observation 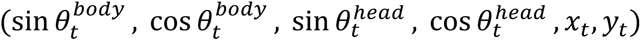 (Figure 1D). Each observation the agent received included these six variables from frames over the past five seconds, making agent inputs 90-dimensional (6 variables × 3 frames per second × 5 seconds, Methods) at each timestep.

**Figure 1.**
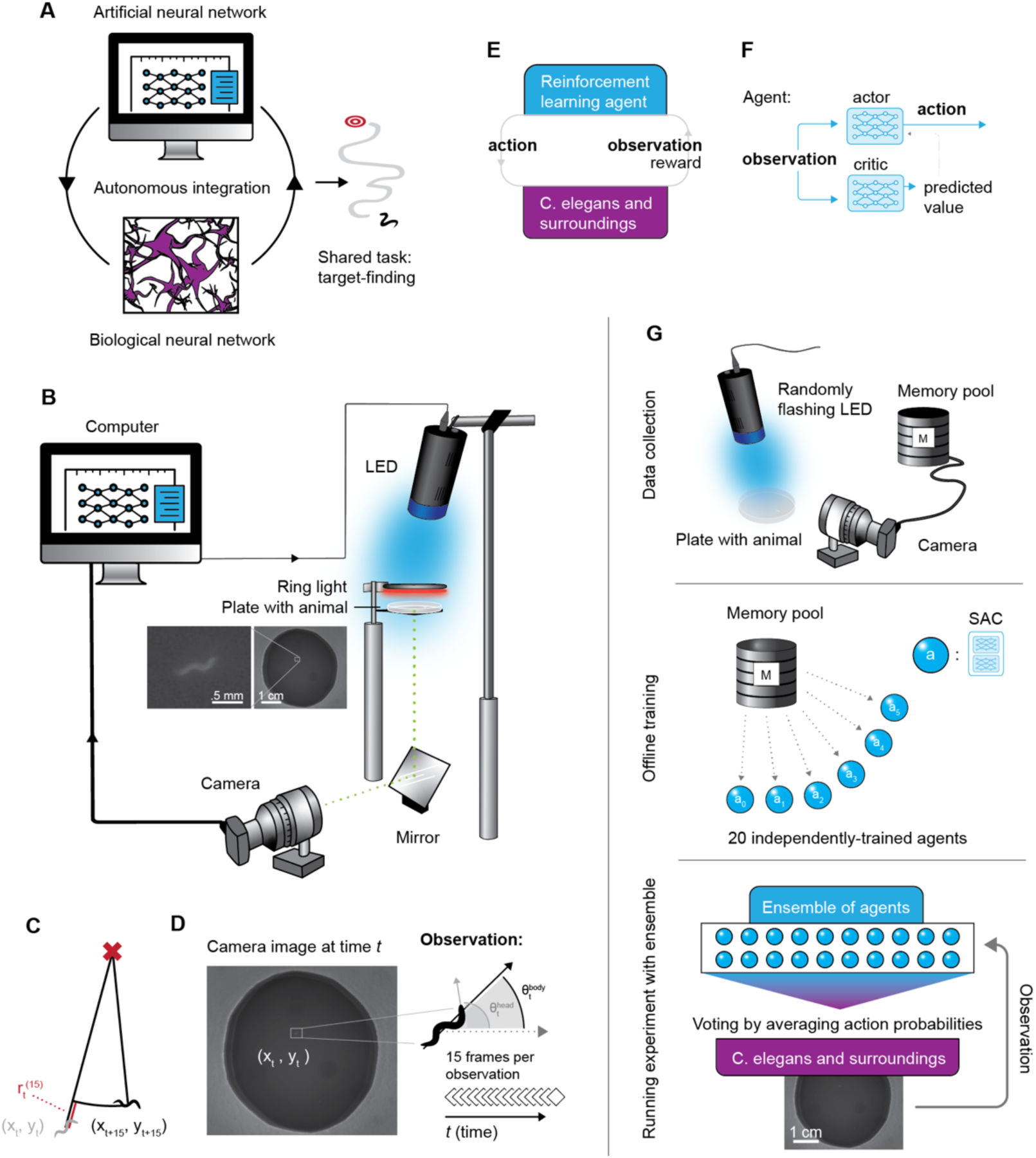
A system that integrates deep RL with the *C. elegans* neural network. (A) Concept for combining artificial and biological neural networks for a shared task. (B) Closed-loop setup using optogenetics. A single nematode was placed in a 4 cm-diameter field and illuminated by a red ring light for imaging. A camera and a high-powered LED (blue or green) were connected to a computer to form a closed-loop system. The LED modulated neurons carrying optogenetic constructs (see main text). (C) Reward at time *t*, *r*_t_ ^(15)^ was defined as the change in distance to target between times *t* and *t+15*. (D) Sample camera image at time *t*. An observation was a stack of 6 measurements from 15 frames (5 s at 3 fps) for a total of 90 variables per observation received by the agent at each timestep. Measurements were coordinates of the animal’s center of mass at time *t* (xt, yt), and the sines and cosines of the head and body angles, 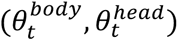 of the animal relative to the positive x-axis. (E) RL loop diagram of the combined system. (F) Actor-critic architecture used as a deep RL agent. (G) Pipeline for training and evaluating the RL-animal system (see main text and Methods for details). A total of 5 h of data were collected where a light is flashed randomly on an animal, stored in a memory pool. Animals were switched out approximately every 20 minutes. Twenty soft actor-critic agents were independently trained on the memory pool. During evaluation, the agents were put into an ensemble that voted in real time on actions. Each individual agent’s decision was based on the observation received from the camera.

Given an observation at time *t*, the RL agent was trained to learn what action *a*_*t*_to take at that time to maximize the return, defined as a sum of rewards discounted over time (Figure 1E, Methods). To take an action, the agent could use optogenetics^14^ to stimulate selected neurons that expressed channelrhodopsin, a light-gated ion channel that can be stimulated by blue light (480 nm) to activate neurons^15^, or archaerhodopsin, which inhibits neurons upon stimulation with green light (540 nm). An agent thus influenced animal behavior by deciding whether to turn an LED on or off at each timestep.

For the implementation of the RL agent, we chose the soft actor-critic (SAC) algorithm because of its successes in simulated and real-world RL environments^27, 31–33^. SAC has separate neural networks for a critic that learns to evaluate observations and an actor that learns to optimize actions based on the critic evaluations and maximize return (Figure 1F, Methods). Both neural networks take observations as input and consist of two layers with 64 units per layer (Methods). The actor outputs probabilities of turning the light on at time *t*, *p*(*a*_*t*_ = 1). We assigned the agent’s action for that observation as “light on” if the actor’s output *p*(*a*_*t*_ = 1) ≥ 0.5.

Deep RL tends to require a large amount of data for training. For instance, agents learning to play Atari can require thousands of hours of gameplay to achieve good performance^23, 24^. It was infeasible to collect thousands of hours of recordings in our environment, and unlike video games or physical systems with reliable dynamics, adequate computer simulations of the *C. elegans* nervous system and its behaviors are not available to generate training data^34^. Therefore, to facilitate algorithm development and reduce the amount of data needed to learn the target-finding task, agents were trained offline on pre-recorded data, which were collected for 20 min per animal for a total of 5 h. During training data collection, the light was turned on randomly with a probability of 0.1 every second (Figure 1G, top and Methods). Following approaches in supervised learning^35^, the data were then augmented during training by randomly translating and rotating the animal in a virtual arena approximately the size of the 4 cm-diameter evaluation arena (Methods).

During training, deep RL agents were unstable and prone to sudden performance drops in the target-finding task (Figure S1), similar to observations from previous work^36, 37^. In simulated environments, such performance crashes can be quickly monitored using evaluation episodes in the exact environment used for testing. In our environment, evaluation episodes were impractical because they would have required many more times the amount of data than were used to train agents. Therefore, we tested several regularization methods to help with stability and found that ensembles of agents were the most effective for our environment (Figure S2-S4). The final deep RL agents were ensembles of 20 SAC agents, and the collection, training, and evaluation pipeline is shown in Figure 1G.

## Agents could navigate animals to targets

As a first step, we used the transgenic line P*ttx-3::ChR2*, referred to as Line 1 in the text (Figure 2A, Table 1). In Line 1, the *ttx-3* promoter drives expression of channelrhodopsin in AIY interneurons, known to be involved in chemotaxis. Prior work has established a deterministic strategy for navigating animals using optogenetic activation of AIY^16^, which we used here as a “human expert” standard to see whether our agent could achieve similar performance.

**Figure 2.**
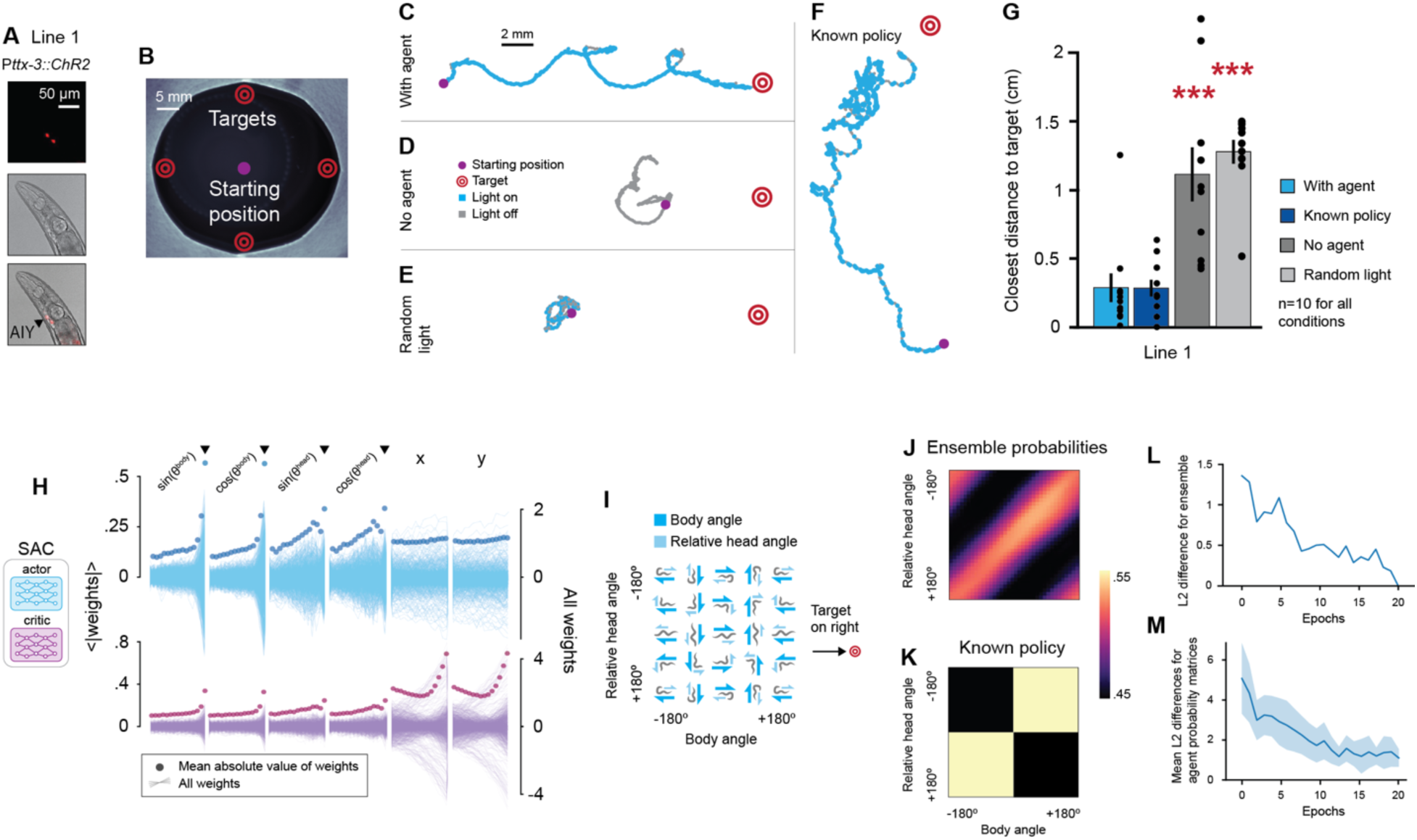
The system learned to navigate the *C. elegans* Line 1 to a target. (A) Optogenetically modified neurons AIY (black arrow) in Line 1. (B) Evaluation setup. The animal was placed in the center (purple circle) of a filter paper circle with diameter 4 cm. In each 10 min episode, agents were tested on their ability to navigate the animal to one of the four target locations shown (red). (C-F) Sample tracks for evaluations with agent, without agent, with random light, and with a “human expert” policy from previous literature (Kocabas et al., 2012), respectively. (G) Closest distance to target achieved by animals for trials with and without an agent as well as with random light stimulations (n=10 for each condition). Animals with agents moved significantly closer to targets than animals without agents. Error bars denote standard error. Mann-Whitney U Test, with agent vs. with control conditions indicated by asterisks, **P<.01, ***P<.001. (H) Weights of the first 64-neuron layer in all actor (top) and critic (bottom) networks in the agent ensemble. For angle-related variables, the most recent frames (black arrows) had the largest weights. (I) Reference for the policy plots in J-K, showing example animal conformations. (J) Trained agent probabilities for simulated inputs. (K) The human expert policy plotted for comparison. It is similar to the learned agent policy, but not identical. (L) The L2 difference in the policy matrix between the final ensemble and ensembles at each epoch during training. By definition, the difference is 0 at epoch 20. (M) Mean L2 differences between individual agents and the final ensemble, with standard deviation shaded in blue.

**Table 1.**
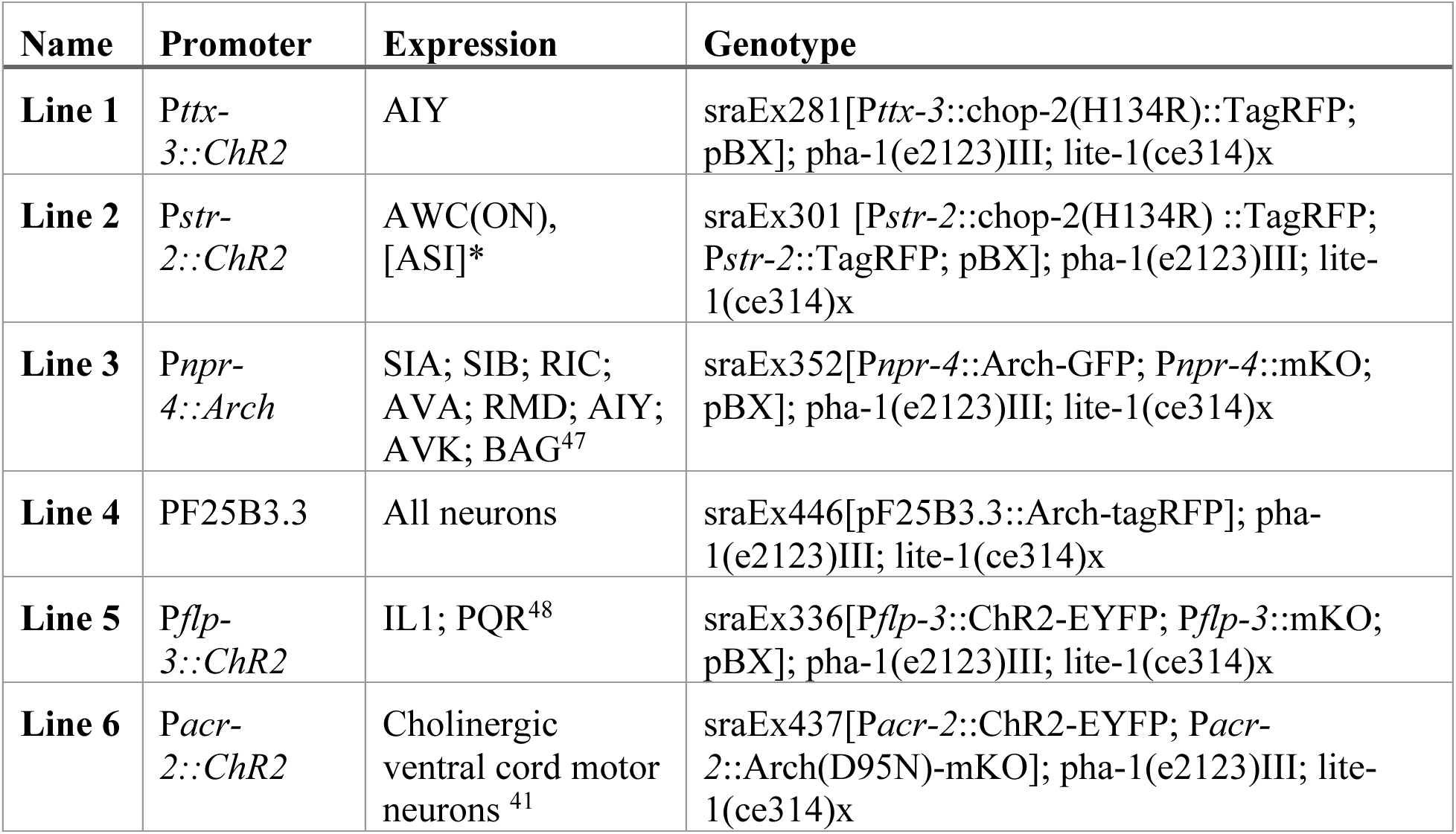
Transgenic line names in text with their genotypes and expression.

After training an RL agent on Line 1, the trained agent was evaluated by placing an animal in the center of a 4 cm-diameter arena and entering target coordinates as input to the agent (Figure 2B). The agent was set to navigate the animal over a 10 min episode to a single target placed in one of four possible locations. The agent learned the pattern in which to turn on the light (blue points) to maneuver the animal toward the target. A sample track of an animal driven by the agent is shown in Figure 2C (see also Video S1). In this example, the animal successfully reaches the target. In contrast, when the animal is disconnected from the agent either by having light off all the time (Figure 2D), or by turning the light on randomly (Figure 2E, Video S2), the animal fails to reach the target. For comparison, we also considered the case where the light was turned on according to a human expert policy, which was successful in driving the animal to the target (Figure 2F). Figure 2G shows statistics for each condition: the closer the distance to the target, the better the performance. The agent’s learned policy performed as well as the human expert known policy, and both of those performed significantly better than controls (learned policy: p<.0006, no agent; p<.0002, random light. Human expert policy: p<.0007, no agent; p<.0002, random light).

To understand what the agent trained on Line 1 learned and compare it to the known policy^16^, we sought a representative subspace of the 90-dimensional observation space in which to plot agent decisions. For every SAC agent in the ensemble, we plotted weights of the first layer of the actor network to assess which input variables were associated with large weights (Figure 2H, S5). Measurements of head and body angles corresponding to the most recent frame in an observation (black arrows in Figure 2H) had larger weight magnitudes than ones from earlier frames. Therefore, to visualize agent strategies, we fixed the values of the 30 coordinate variables ((*x*_*t*)_, *y*_*t*)_); *t* − 5 *s* < *t*’ < *t*) in each observation to a position left of the target (Figure 2I, Methods) and plotted the probability that the ensemble turned the light on as a function of body and head angles at the latest time in the observation 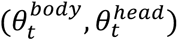 (Figure 2J). Likewise, the human expert policy was plotted in Figure 2K.

To interpret the policies, it is useful to compare Figures 2I and J. The diagonal band in Figure 2J corresponds to the same diagonal in Figure 2I where the animal’s head is always pointed toward the target. There was a high probability of turning the light on along the diagonal, *i.e.,* when the animal’s head was aligned toward the target. Interestingly, the agent’s learned policy was conceptually similar but quantitatively slightly different from the known expert policy plotted in the same projection in Figure 2K, which had a greater emphasis on turning animals in the correct direction. Nonetheless, both policies were effective in this targeted navigation task.

The projection in Figure 2J also provided a way to plot agent training progress over time, with Figure 2L-M showing the change in agent policies over 20 epochs of training. Figure 2L is the difference between the policy of full ensembles during training and after training, while Figure 2M takes differences between individual agent policies and compares them to the final trained ensemble, plotting average differences with standard deviations. The range in y-axes in these figures shows that individual agents, even after training, can be quite far from the final policy compared to ensembled ones during training, highlighting the importance of an ensemble of agents.

## The agent learned policies based on its site of integration

We aimed to build a robust and flexible algorithm that could be trained to adapt to its connected neurons. We next tested whether the RL agent could learn appropriate rules for a variety of neural connections without any explicit prior knowledge about them. New agents were trained on five additional transgenic lines that were functionally distinct from Line 1 and did not have associated human expert policies as benchmarks (Figure 3). These lines are ordered in the text by agent performance in navigating animals to targets compared to no light and random matched-frequency light controls. See Table 1 and Figure 3A for line genotypes and neuron expression.

**Figure 3.**
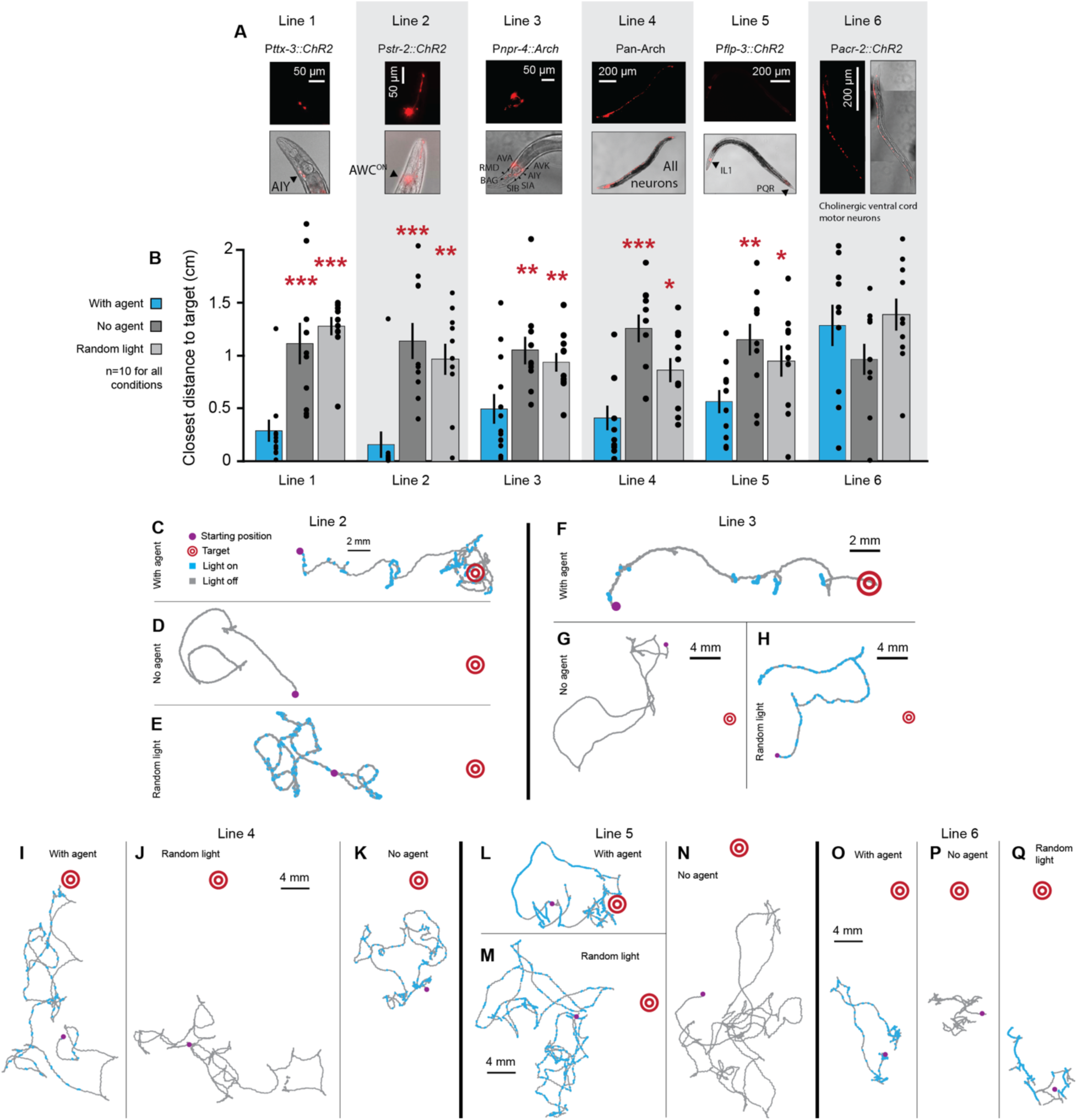
The system could successfully navigate different optogenetic lines to a target. (A) Images of optogenetic lines with promoters and modified neurons. (B) Statistics for each line (n=10) comparing performance with agents, without agents, and with frequency-matched random light controls. Error bars denote standard error. Mann-Whitney U Test, with agent vs. with control conditions indicated by asterisks, **P<.01, ***P<.001. Lines 1-5 were successful. (C-E) Following the format in Figure 2C-F, example tracks for Line 2 with positions of light activation along the trajectory highlighted in blue for animals C, with the agent, D, without any optogenetic activation, and E, with randomly flashing light. (F-Q) Example tracks for Lines 3-6 for each experimental condition in C. Variability in starting positions for controls can be explained by free movement in the time between placing animals on the plate and starting the experiment, approximately 1 min.

Lines 3-6 expressed light-sensitive channels in multiple neuron types. Line 3 and 4 animals, unlike the other lines that expressed channelrhodopsin, expressed archaerhodopsin, which inhibits neurons upon stimulation with green light (540 nm). For Line 4, due to concerns about phototoxicity, agents were restricted to pulses of light during evaluation (Methods). These lines tested the abilities of the RL agent with different sets of neuronal connections and different means of neural modulation.

In Lines 1-5, animals with trained agents moved closer to targets than control animals did (Figure 3B). Example tracks showing agent activity during evaluation and controls are shown in Figure 3C-Q. Videos S1-6 show agent performance and controls for Lines 1-3, which performed best of the six lines. Given that policies for goal-directed movement using optogenetic modulation of these lines were previously unknown, it was remarkable that agents still learned to direct these animals towards a target (for Line 3, see Bhardwaj et al., 2018^38^ for *npr-4* mutant behavior and for Line 5, see ^39^ for IL1 involvement in head withdrawal).

The agent was able to successfully interact with Lines 3-5, each of which involved multiple neurons (Figure 3B). It was surprising that agents learned a policy for Line 4 in which they were coupled to the entire nervous system^40^. In this instance, the agents took advantage of increased movement after a period of freezing, in contrast to the Line 3 policy that relied on slowed movement or turning during neuron inhibition. However, the agent failed to find an effective policy for Line 6, in which it was coupled to cholinergic muscle excitation in the ventral cord^41^. The standard deviation in the learned policy between agents in the ensembles was noticeably greater for Lines 4-6 (Figure 4A-B), which had poorer performance than Lines 1-3 (Figure 3B). These results together show that the site of integration plays an important role in the success of the animal and agent system, and that not all neurons or circuits can be modulated to drive an animal to a given target.

**Figure 4.**
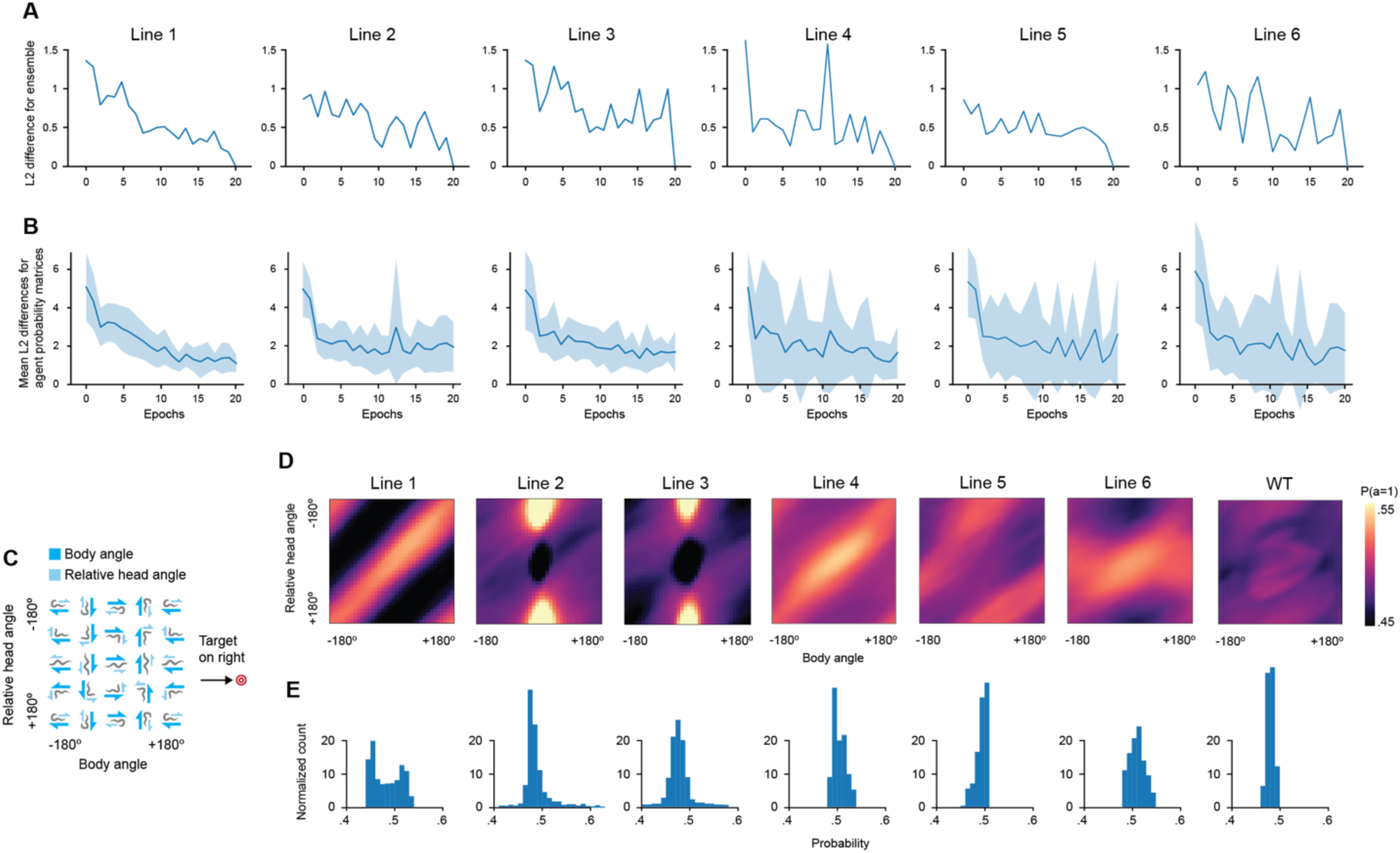
The system learned to navigate different optogenetic lines to a target with neuron-specific strategies. (A) L2 differences between final ensembles and ensembles at each epoch during training. (B) Mean L2 differences between individual agents in the ensemble and the final trained ensemble. Within an ensemble, agents for Lines 4-6 varied more than in Lines 1-3, which is reflected in the narrower range of probabilities for Line 4-6 in B. (C) The animal conformation reference plot for agent policies in B (repeated from Figure 2I). (D) All agent policies for Lines 1-6, and an agent trained on wild type data where there was no possible successful policy. Lines 1 and 4, as well as 2 and 3, had similar agent policies. (E) Probabilities in D plotted as a histogram. Lines 1-3 had larger ranges, suggesting greater certainty.

We visualized the resulting policies using the same metrics from Figure 2I-J to understand how the interfaced neurons were involved in navigation toward the target. Figure 4C again shows animal postures used in mapping out agent policies for reference. The policies are plotted in Figure 4D. Ensemble action certainty is also visible in Figure 4D-E, in which Lines 1-3 have probability values with a wider range than Lines 4-6. This indicates that agents are more confident about when to turn the light on or off in Lines 1-3. For comparison, we show an agent trained on wild type animals (Figure 4D), which had no response to optogenetic modulation. The policies described in Figure 4D show that agents learned strategies tailored to the neurons with which they were interfaced.

## Agents predicted similarities between neural circuits

Broadly, there were three successful strategies represented by Lines 1 and 4, Lines 2 and 3, and Line 5 (Figure 4D). To understand how the agent policies interacted with the nervous system, we focused on the three most successful lines: Lines 1, 2, and 3. Although the behavior of Line 1 in response to blue light is mostly to move forward and the behavior of Line 2 is mostly to reverse, agent policies were not merely inverses of each other. Rather, agents learned that Line 1 control was dependent largely on the animal’s head angle relative to the target while Line 2 and 3 control depended on specific head and body angle combinations. Despite large differences in Lines 2 and 3 (excitation of a single neuron in Line 2 versus inhibition of multiple neurons in Line 3), training on Line 3 resulted in an action probability matrix that was strikingly similar to the one from training on Line 2.

To quantify these similarities in learned actions for the different lines and to assess the degree of generalization across different sites of integration, we ran target navigation experiments where each agent was tested on each of these three lines (Figure 5A). Sample tracks from combinations of agents and animals are shown in Figure 5B, with average results in Figure 5C. To evaluate whether agent policies were predictive of cross-evaluation performance, we measured L2 norm differences of the action probability matrices (Figure 5D). This is a symmetric matrix that reveals the differences between the different policies. As intuitively observed in Figure 5D, the policies from Lines 2 and 3 are most similar. The corresponding plot using the experimental data from Figure 5C is shown in Figure 5E. As expected, diagonal entries reveal low distance to targets. Line 3 animals tested with Line 2 agents also showed low distance to targets.

**Figure 5.**
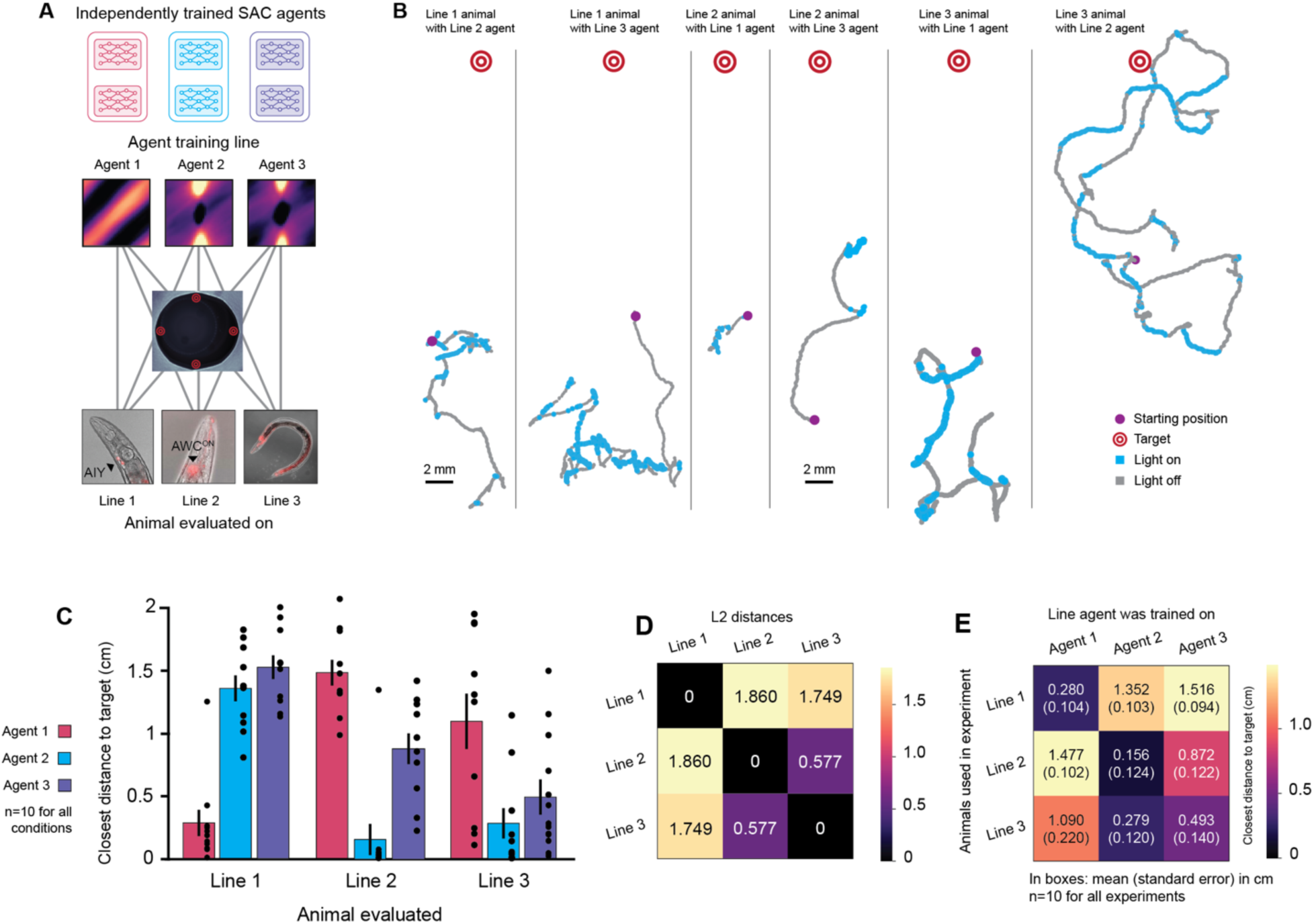
Agent policies can predict agent performance on other lines. (A) An illustration of cross-evaluation experiments, in which agents trained on each of the 3 best-performing lines were evaluated on every other line. (B) Sample tracks with agent actions for each combination of agent and animal not shown in Figure 2C, 3C, or 3F. (C) Statistics of closest distance to target for each combination of agent and animal with n=10 per condition. (D) L2 distances between ensemble action probability matrices for each genetic line. (E) Mean closest distances (cm) to the target in a 10-min evaluation episode is shown with standard error in parentheses. Distances between the ensemble action probability matrices in D correlate with the closest distances achieved in across-policy evaluation experiments (r2=.8578, p <.0004).

The matrix of cross-evaluation results in Figure 5E correlated well with predictions based on the similarity of the action probability matrices in Figure 5D (r^2^=.8578, p <.0004). As expected from the contrast in action probabilities in Figure 4D, Line 1 versus Lines 2 and 3, Line 1 did not respond well to agents trained on Lines 2 or 3. For example, when the agent trained on Line 1 was tested with an animal from Line 2, the closest distance reached from the target was about 1.477±0.102 cm, much larger than when tested on Line 1, 0.280±0.104 cm (Figure 5E). The closest distance was also comparable to or greater than the no agent or random light conditions for Line 2 (Figure 3B), as the Line 1 agent tended to drive Line 2 animals away from rather than toward targets (p-value<.08, no agent; p-value<.009, random light; Mann-Whitney U Test). Likewise, neither Line 2 nor 3 animals performed well on the task when paired with the Line 1 agent. In summary, by comparing action probabilities learned by agents that were trained to couple to specific sets of neurons, we could make accurate predictions about the behavior of these lines under optogenetic control in the target-finding task.

Another particularly interesting finding was that Line 2 and 3 animals were most successful when paired with the Line 2 agent even though the Line 3 agent was trained on data from the line itself (p<.002, Line 2 line with Line 2 vs. Line 3 agent; p<.04, Line 3 line with Line 2 vs. Line 3 agent, Mann-Whitney U Test, n=10). These results may be explained by higher data quality caused by the stronger response of Line 2 to optogenetic stimulation (Supplementary Videos 1, 2, 5, 6), reflected in the greater action certainties in the Line 2 ensemble as compared to the Line 3 ensemble (Figure 4D). This result suggests that training RL agents with less action noise could improve performance in noisy biological environments^42^. Overall we demonstrate that our system may be used to generate hypotheses about learning in biological environments with greater access to internal mechanisms (through the artificial network) than an animal’s nervous system alone can provide.

## Animals corrected errors made by agents during food search

We next evaluated whether agents and animals could transfer their abilities from the target-finding task to improve food search. Using the three best-performing lines, we tested whether the animal could correct errors made by an agent about the location of food. For these error-handling tasks, targets for the AI agent were placed at increasing distances from the edge of the actual 5 µL patch of food (OP50 *E. coli* bacteria) to mimic errors made by the agent (Figure 6A; Methods). Agents were on throughout the experiment: crucially, the agents remained active even after animals reached the target. Animals were tested on whether they could reach the food in 20 min trials with or without RL agents. Agents were identical to the ones used in Figure 2-5, with each line tested using its own agent. For both Lines 1 and 2, when targets were 0.5 cm away from food edges, animals were able to leave an agent’s target region (a circle of radius 0.0625 cm; Methods) and moved to the food in 8/10 trials (p<.0004) (Figure 5B-C). This was significantly different from trials without any agent assistance, in which 0 animals reached food in 10 trials for both lines. Line 3 was not as successful with agent assistance (Figure 6D), likely due to the less reliable control in moving animals to a target (Figure 3B). This suggests that simultaneous modulation of the neurons in this line is not as strongly linked to directed movement as in Lines 1 and 2. In contrast, Line 1 and 2 animals could effectively switch between making decisions based on their own sensory systems or the agents, which were trained to keep animals at targets. Sample tracks for all experimental conditions are shown in Figure 6E-G.

**Figure 6.**
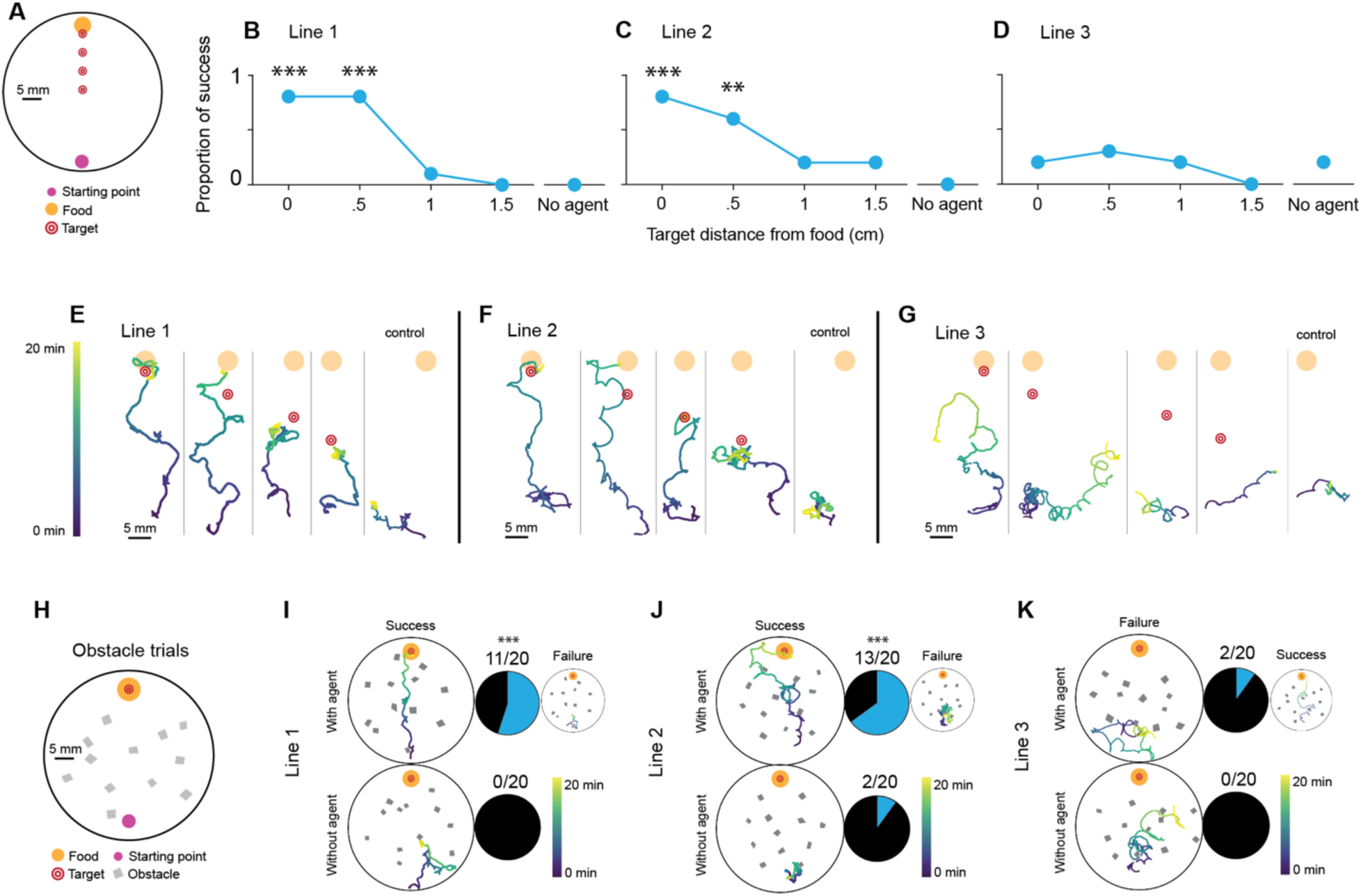
Animals with agents can correct errors and generalize to novel situations. (A) Diagram of error-handling food search experiments. A single animal was placed at the opposite end of a plate (starting location large purple circle) as a 5 µm drop of OP50 *E. coli* bacteria (orange circle). Trials lasted 20 min and success was defined by whether the animal reached food. Agents were directed to navigate animals to a target a distance away from the food (agent target location denoted by concentric red circles). (B-D) Proportion of animals that successfully reached food for Lines 1-3, respectively, n=10 for each condition. For Lines 1 and 2, targets up to 0.5 cm away led to significantly better performance than without agents. **P<.01, ***P<.001 (with agent vs. no agent; p<.0004 for Line 1 with target at 0 and 0.5 cm from food and Line 2 with target at 0 cm from food; p<.006 for Line 2 with target at 0.5 cm from food), (E-G) Sample tracks for Line 1-3 animals with agents based on the majority result of trials with the target at the given distance. A control track without an agent is shown in the fifth column. (H) A diagram of the plate used for experiments with obstacles. Twelve paper rectangles with side lengths approximately 2 mm were scattered on the plate. Agents were directed to navigate animals to the food and again, success was determined by whether animals reached food. (I) Sample tracks for Line 1 animals that succeeded (top left) or failed (top right), with control trials without agents (bottom). Success rates shown in pie charts. Animals with agents were significantly more likely to succeed; ***P<.001 (permutation test). (J) Sample tracks for Line 2 animals. 13/20 animals succeeded with agents and 2/20 without. Animals with agents were significantly more successful. (K) Sample tracks with Line 3 animals, with a failed trial in the top left to represent the majority outcome. 2/20 animals reached food with agents and 0/20 without (permutation test, p=.244).

## RL agents with animals could navigate novel unforeseen environments

To test the generalization abilities of the RL agent/animal system, we next tested whether the animal and agent could navigate an unforeseen environment with obstacles to reach food. This scenario represented a novel environmental condition that neither the agent nor the animal had seen before. We designed trials in which twelve paper quadrilaterals with 1-3 mm edges (comparable to the 1 mm body length of *C. elegans*) were scattered randomly on the plate to serve as obstacles between an animal and a 5 µL patch of food (Figure 6H; Methods). In this scenario, animals were again tested on whether they could reach food during a 20 min trial with and without agents. This was a particularly challenging task because animals had to use their sensory and motor systems to navigate around obstacles, while agents had to navigate animals to food despite noisy movements caused by obstacles. Line 1 and 2 animals performed very well in navigating this new environment to find food (Figure 6I-J, p-value<.0001, Line 1; p-value<.0004, Line 2; permutation tests). Line 3 was not as successful (Figure 6K); overall, the agent could navigate Line 3 animals closer to targets but could not achieve more difficult food search tasks. For Lines 1 and 2, however, these data provide evidence that our system displays cooperative computation between artificial and biological neural networks to improve *C. elegans* food search in a zero-shot fashion without any retraining in novel environments.

### Discussion

We showed here how to build a hybrid system where deep RL can interact with an animal’s nervous system to improve a target behavior. In the data-limited context of biological systems, we could train deep RL agents using data augmentation and improve the stability of deep RL using an ensemble of agents. Agents could customize themselves to specific and diverse sites of neural integration. These results did not depend on the number of neurons that agents were interfaced with, nor whether the interactions were excitatory or inhibitory. A failure to learn, as in Line 6, shows the critical importance of the neural circuit under control and demonstrates that not all circuits can be successfully modulated by our RL agents. In addition, the animal plus agent system could generalize a learned target-finding strategy to novel environments for food search. We demonstrated that the inherent ability of the *C. elegans* nervous system to find food could be enhanced by deep RL, helping animals find targets faster and in more challenging environments than they could on their own.

In previous work, brain-machine interfaces have allowed animals to control machines through neural recordings^43–45^. Conversely, supervised optogenetic manipulations have taken control of *C. elegans* neurons or muscles to turn the animal into a passive robot^16, 46^. In contrast to both of these types of artificial-biological neural interactions, our work integrated a living nervous system with an artificial neural network, automatically discovered activation patterns to interact with the nervous system, and did so in a way that allowed computations from both networks to drive animal behavior in a robust manner that generalizes in a zero-shot fashion to novel environments. Our system was also able to discover patterns of neural activity that were sufficient to drive specific behaviors: studies of sufficiency complement the more traditional lesion and inhibition studies in neuroscience, which have historically only focused on determining the neural circuitry correlated with or necessary for specific behaviors.

We used *C. elegans* as a model organism for its small and accessible nervous system. It would be interesting for future work to test our method in larger state spaces and action spaces, as one would find in an animal with a richer behavioral repertoire and larger nervous system. Deep RL has already solved complex simulated tasks in high dimensional spaces with large numbers of parameters^21, 23, 25^, suggesting its potential for integration with larger animals. Overall, our study opens new avenues for understanding neural circuits, improving behavior using deep RL, and building hybrids between artificial and biological networks that can utilize the flexibility, robustness and computational power of AI and animals.

### Methods

## Animal genetics and care

### Genetic lines

Strains are listed in Table 1. All animals had *lite-1* mutant backgrounds to reduce light sensitivity. Lines were chosen after an initial screen for response to optogenetic activation or inhibition.

### Animal maintenance

1. *C. elegans* strains were cultured at 20°C (room temperature) on nematode growth media (NGM) plates seeded with *E. coli* strain OP50. Animals used in optogenetic experiments were cultured at 20°C on NGM plates seeded with *E. coli* strain OP50 with 1 mM all-trans-retinal (ATR) at a 9:1 volume ratio, for at least 12 h before experiments. (ATR is a cofactor required for rhodopsin activity.)

## Experimental setup

### Experimental system hardware

Experiments were conducted at 20°C. Two setups were built as in the diagram in Figure 1b. The first used an Edmund Optics 5012 LE Monochrome USB 3.0 Lite Edition camera. The assay plate was lit with an Advanced Illumination RL1660 ring light. For the second rig, the camera was a USB-connected ThorLabs DCC1545M. Both cameras were run at 3 fps, which was a rate slow enough for image capture, image processing, action decision, and action transmission to occur. Lights for optogenetic illumination were Kessil PR160L LEDs at wavelengths of 467 nm for blue and 525 nm for green. The plate was illuminated with a Grandview COB Angel Eyes 110mm Halo ring light. Kessil LEDs for optogenetic activation were controlled by a National Instruments DAQmx that was in turn managed through a Python library.

### Animal tracking

For all experiments animals were moved from food plates to a 10 cm-diameter NGM tracking plate. Tracking plate setups depended on the experiment, but all plates had a filter paper ring to confine the animal to a 4 cm-diameter circle. We soaked the paper in 20 mM copper (II) chloride solution, an aversive substance to *C. elegans* before placing it on the plates. Obstacles used in Figure 6h-k were not soaked in copper solution. If food patches were used in the experiment as in Figure 6, 5 µL of OP50 *E. coli* bacteria were deposited on the plate and allowed to grow at room temperature (20°C) for roughly 24 hours.

## Collecting training data

Five hours of data were collected for each genetic line in 20 min episodes. In every episode, a single nematode cultured with ATR was placed on an NGM plate. As in the animal tracking setup, a filter paper barrier of diameter 4 cm was placed on the plate. A camera then recorded images at 3 fps while a blue or green LED flashed randomly on the plate. Blue light was used for animals modified with channelrhodopsin and green light was used for animals modified with archaerhodopsin. A decision to turn the light on or off was made every 1 s with a probability of 10% on. If on, the light duration was also 1s. Animals were switched out for new ones after each episode. Light decisions and images were stored for agent training in separate datasets for each line.

## Reinforcement learning details

Reinforcement learning (RL) is a framework in which an agent interacts with an environment and attempts to maximize a reward signal. The agent receives observations from the environment, giving it an idea of the environment’s current state, and learns what actions to take that will be most likely to maximize the reward signal received from the environment. The RL agent learns through experience an action probability distribution, π(*a*_*t*_|***s***_***t***_), where *a*_*t*_is the action taken at time *t*, ***s***_***t***_ is the state received from the environment corresponding to time *t*, and the maximized reward *r*_*t*_ is received at time *t*. Each of these variables is defined below.

We used a discrete soft actor-critic (SAC) algorithm for all agents^31, 32^. For each genetic line, 20 SAC agents were independently trained offline on the same data pool.

### Variable definitions

#### Observations

Every camera image was preprocessed into features known to be relevant in *C. elegans* behavior^16^. We used pixel coordinates (*x*, *y*) of the animal’s centroid location in the image, the body angle relative to the +*x*-axis and the head angle relative to the +*x*-axis (see Figure 1). Body angles were computed by fitting a line to a skeletonized worm image and head angles were computed through template matching. See the code in `improc_v.py` for details.

Head/tail identification was done by assigning the head label to the endpoint that was closest to the head endpoint in a previous frame. To handle reversals, a common behavior in freely moving animals, the overall movement vector over 10 s was compared to tail-to-head vectors during the same window of time. If the vectors pointed in different directions, head and tail labels were switched. Before each evaluation episode, 5 s of frames were collected to assign the first head label again by comparing movement vectors to tail-to-head vectors.

Angles were converted to sine and cosine pairs to avoid angle wraparound issues. 15 frames (5 s at 3 fps) were concatenated together for a single observation. Coordinates were normalized so their means in each 15-frame observation was within [−0.5, 0.5]. An observation ***s***_***t***_corresponding to time *t* was thus comprised of 6 × 15 = 90 variables:

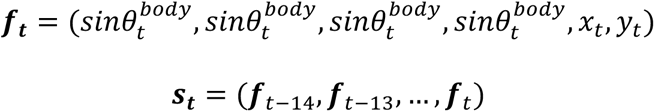

Above, ***f***_*t*_ denotes the tuple of variables for the frame at time *t*. See Figure 1D for a diagram defining the head and body angles.

#### Actions

An action at time *t*, *a*_*t*_, was defined as a choice between the options “light on” or “light off,” denoted by a binary 0 or 1 signal.

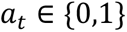

We did not place any constraints on actions, as all ensembles learned policies with overall light exposure that was under 50% of the time (see Methods: Standard evaluation).

#### Rewards

Reward *r*_*t*_ was based on the target-finding task and defined as the distance moved toward the target between the time of the action *t* and 15 frames (5 s) after the action (Figure 1C).

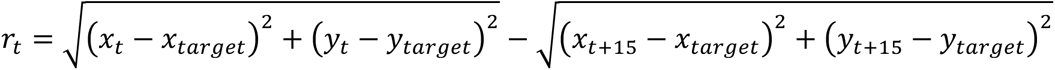

A target region was defined as a circle of radius 30 pixels (625 µm). If the animal was within the target region, the calculated reward was replaced by a constant reward of 2. All other rewards were scaled by a factor of 2 to normalize values and facilitate training.

### Training

As in standard reinforcement learning, SAC searches for a policy π(*a*_*t*_|***s***_***t***_) for an environment with a transition distribution *ρ*_*π*_. π(*a*_*t*_|***s***_***t***_) is the probability of taking an action *a*_*t*_ given an observation ***s***_***t***_. Here we also make explicit the dependence of *r*_*t*_on ***s***_***t***_and *a*_*t*_. SAC deviates from the standard goal of maximizing the return, or expected sum of rewards over time,

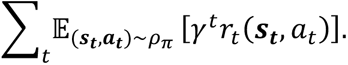

Here, *γ* (fixed at 0.95) is a temporal discount factor that diminishes rewards far into the future. SAC maximizes not only the expected sum of rewards, but also an entropy term weighted by a temperature parameter *α*:

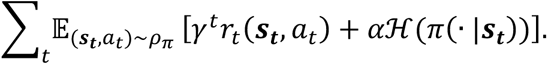

The sum now contains an added entropy term ℋ of the policy π(·|***s***_***t***_), scaled by a temperature parameter *α*.π( |***s***_***t***_) signifies the policy function *π* over all possible events. We used a discrete version of SAC with automatic entropy tuning (see code for implementation).

#### Data augmentation

Once data were collected, they were stored in a memory buffer as tuples:

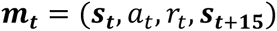

At each training step, a batch of 64 memory tuples were randomly drawn from the buffer and independently augmented by a random translation and rotation. First, the tuple was centered such that the average of the location coordinates were at the origin, (0,0) pixels. Then a location within a ±450-pixel square (comparable to the size of the evaluation arena) was drawn from a uniform distribution and the coordinates recentered around that location. An angle was likewise chosen from a uniform distribution [0°, 360°) and added to the measured angles in the memory tuple.

#### Training details

See Table S1 for architecture and hyperparameter choices. 20 agents per genetic line were trained independently on the same memory buffer for 20 epochs of 5000 steps each. Minibatch size was 64. Weights were initialized using Xavier uniform initialization and biases were initialized at 0. We tried dropout and weight decay on actors, critics, or both, and found that none of these regularizers helped enough to compensate for the need to choose more hyperparameters (see Figures S2-S4).

Independent agents were trained such that the randomly taken action *a*_*t*_, reward *r*_*t*_, and the associated states ***s***_***t***_ and ***s***_***t* * 15**_ were used to learn a state-action value function. This is called a Q-function and was learned by the critic network. The actor network then learned a policy that was the exponential of the Q-function. See Haarnoja et al., 2018^31^ for details.

#### Ensembles

Once the 20 agents for one ensemble were trained, they were combined by taking the average of their action probabilities and setting a threshold at 0.5. That is,

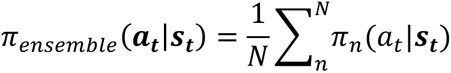

where *N* = 20. If the average probability *π*_ensemble_(*a*_*t*_|***s***_***t***_) ≥ 0.5, then the light was on at that timestep. 3-5 random seeds were run for each genetic line, and the final ensemble was chosen based on inspection of visualized agent strategies.

### Compute resources

All training was done on the FASRC Cannon cluster supported by the FAS Division of Science Research Computing Group at Harvard University. Every agent was trained on a compute node with one of the GPUs available on the cluster: Nvidia TitanX, K20m, K40m, K80, P100, A40, V100, or A100.

### Agent strategy visualization

To visualize agent decisions, we simulated animal states in a smaller space than the full 90-dimensional inputs based on input weight magnitudes. Because the final timesteps of all angle measurements had larger magnitudes than previous timesteps (Figure 2H, Figure S7), we chose to keep input angles constant within each observation and explored the full range of angle possibilities [−180°, 180°) in increments of 10° for 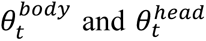 (36 values each). The 30 coordinate variables (*x*_*t*)_, *y*_*t*)_); *t*’ − 5 < *t*’ < *t*) were always fixed to 0.94 cm to the left of the target, which was exactly half the maximum distance used for random translations during training. In total, 36 head angle values × 36 body angle values gave rise to 1296 different input observations, each of which were given to an agent ensemble that then output the decision probabilities recorded in the resultant action probability matrix.

## Evaluation

All experiments involved a single animal placed on a 10 cm-diameter NGM plate with a 4cm-diameter filter paper barrier soaked in copper (II) chloride. All animals were cultured on food with ATR and were thus sensitive to optogenetic perturbation.

### Standard evaluation

Animals were placed in the center of the field. A target was randomly chosen among top, bottom, left, and right options (Figure 2B). The experiment with agents were run for 10 minutes each at 3 fps. At the end of the experiment, animals were switched out.

For controls without the agent, animals freely moved on the plate and were recorded for 10 min. A random target was assigned to compare controls to trials with agents.

For controls with random light exposure, the idea was to make sure that light exposure alone was not responsible for more movement, which could lead to an increased rate of success. Once all trials with agents had been run, the proportion of time where the light was on was calculated for each genetic line. These proportions were 0.4647 for Line 1, 0.2896 for Line 2, and 0.3844 for Line 3. Animals were recorded while light decisions were made every 1 s, with the probability of light on according to the genetic lines listed.

For Line 4 (Pan-Arch), due to concerns about phototoxicity, the evaluation was restricted to 1 s light pulses with 4 s rest periods between them.

### Cross-agent evaluation

In Figure 5, trained ensembles of agents were tested on the genetic lines they had not been trained on. The experiments were conducted identically to standard target-finding evaluations. 10 trials of 10 min each were performed for every agent-genetic line combination.

### Error-handling food search experiments

For the food search experiments in Figure 6a-g, a 10 cm NGM plate was prepared with a 4 cm-diameter filter paper circle soaked in 20 mM copper (II) chloride. 5 µL of OP50 bacteria were grown for ∼24 h before experiments.

Each trial lasted 20 min. An animal was placed on one end of the plate with the OP50 droplet at the opposite end. During the 20 min, the same agents trained on random data as in the standard evaluations were set to navigate animals to targets at 0 cm, 0.5 cm, 1 cm, or 1.5 cm away from the edge of the OP50 droplet. For control trials, agents were left off and the animal roamed freely for 20 min.

Success was defined as a binary outcome as in the obstacle experiments. If an animal reached the food within the 20 min trial, it was counted as a success. Out of 270 trials run across all genetic lines involving OP50 droplets (obstacles and food search), only 1 CH1 animal left food after reaching it during a food search trial when the target was placed 1 cm away from the food edge. This trial was counted as a success.

### Obstacle food search experiments

For the obstacle trials in Figure 6H-K, a 10 cm NGM plate was prepared with a 4 cm-diameter filter paper ring soaked in a 20 mM copper (II) chloride solution. We cut 12 pieces of filter paper into quadrilaterals with side lengths 1-3 mm and scattered them on the plate (they were not soaked in copper (II) chloride solution). Sample arrangements are shown in Figure 6H-K. Plates were replaced with new obstacle arrangements every 5-10 trials. 5 µL of OP50 bacteria were grown on one side of the plate for ∼24 h before experiments.

Each obstacle experiment was a 20 min trial. A single animal was placed on one end of the plate as in Figure 6H, with the food droplet on the other end and the obstacles in between animal and food. Trained agents (the same agent ensembles used in standard evaluations) were run on the genetic line they were trained on for 20 min. Agents were not retrained to handle obstacles. Control trials had no optogenetic manipulation; that is, the animal was allowed to freely roam the plate with obstacles and food for 20 min. Success was defined as a binary outcome, indicating whether an animal reached food during the trial.

## Data and code availability

Processed animal tracks, analysis code, and training code examples are available at https://tinyurl.com/RLWorms. Other data are available upon request.

## Author Contributions

CL, GK, and SR designed the study. CL wrote code, performed experiments, and did data analysis. CL, GK, and SR wrote the manuscript.

## Acknowledgments

We thank Surya Bhupatiraju for discussions about reinforcement learning and comments on the manuscript. We thank Timothy Hallacy and Abdullah Yonar for guidance in *C. elegans* experiments and Cory McCartan for input on statistical analyses. We thank Kenneth Blum, Cengiz Pehlevan, Giri Anand, Alexandru Bacanu, Benjamin Brissette, Dianna Hidalgo, Roya Huang, Heitor Megale, William Weiter, Yusuf Ilker Yaman, Vincent Zhuang, and Steven Zwick for comments on the manuscript.

This work was supported in part by NIGMS grant 1R01NS117908-01 (SR), Dean’s Competitive Fund from Harvard University (SR, CL), NIH R01EY026025 (GK), and an NSF GFRP fellowship (CL).

## Competing interests

The authors declare no competing interests.

## Notes

https://tinyurl.com/RLWorms

### Competing Interest Statement

The authors have declared no competing interest.

### Summary of Updates

Additional data on new transgenic lines.

## References

1. LeCun, Y., Bengio, Y. & Hinton, G. Deep learning. Nature 521, 436–444 (2015).

2. Sinz, F. H., Pitkow, X., Reimer, J., Bethge, M. & Tolias, A. S. Engineering a Less Artificial Intelligence. Neuron 103, 967–979 (2019).

3. Afraz, S.-R., Kiani, R. & Esteky, H. Microstimulation of inferotemporal cortex influences face categorization. Nature 442, 692–695 (2006).

4. Bonizzato, M. & Martinez, M. An intracortical neuroprosthesis immediately alleviates walking deficits and improves recovery of leg control after spinal cord injury. Sci. Transl. Med. 13, eabb4422 (2021).

5. Enriquez-Geppert, S., Huster, R. J. & Herrmann, C. S. Boosting brain functions: Improving executive functions with behavioral training, neurostimulation, and neurofeedback. Int. J. Psychophysiol. 88, 1–16 (2013).

6. Iturrate, I., Pereira, M. & Millán, J. del R. Closed-loop electrical neurostimulation: Challenges and opportunities. Curr. Opin. Biomed. Eng. 8, 28–37 (2018).

7. Lafer-Sousa, R. et al. Behavioral detectability of optogenetic stimulation of inferior temporal cortex varies with the size of concurrently viewed objects. Curr. Res. Neurobiol. 4, 100063 (2023).

8. Lu, Y. et al. Optogenetically induced spatiotemporal gamma oscillations and neuronal spiking activity in primate motor cortex. J. Neurophysiol. 113, 3574–3587 (2015).

9. Salzman, D., C., Britten, K. H. & Newsome, W. T. Cortical microstimulation influences perceptual judgements of motion direction. Nature 346, 174–177 (1990).

10. Schild, L. C. & Glauser, D. A. Dual Color Neural Activation and Behavior Control with Chrimson and CoChR in Caenorhabditis elegans. Genetics 200, 1029–1034 (2015).

11. Xu, J. et al. Thalamic Stimulation Improves Postictal Cortical Arousal and Behavior. J. Neurosci. 40, 7343–7354 (2020).

12. Bergmann, E., Gofman, X., Kavushansky, A. & Kahn, I. Individual variability in functional connectivity architecture of the mouse brain. *Commun*. Biol. 3, 1–10 (2020).

13. Mueller, S. et al. Individual Variability in Functional Connectivity Architecture of the Human Brain. Neuron 77, 586–595 (2013).

14. Husson, S. J., Gottschalk, A. & Leifer, A. M. Optogenetic manipulation of neural activity in C. elegans: from synapse to circuits and behaviour. Biol. Cell 105, 235–250 (2013).

15. Nagel, G. et al. Channelrhodopsin-2, a directly light-gated cation-selective membrane channel. PNAS 100, 13940–13945 (2003).

16. Kocabas, A., Shen, C.-H., Guo, Z. V. & Ramanathan, S. Controlling interneuron activity in Caenorhabditis elegans to evoke chemotactic behaviour. Nature 490, 273–277 (2012).

17. Leifer, A. M., Fang-Yen, C., Gershow, M., Alkema, M. J. & Samuel, A. D. T. Optogenetic manipulation of neural activity in freely moving Caenorhabditis elegans. Nat. Methods 8, 147–152 (2011).

18. Wen, Q. et al. Proprioceptive Coupling within Motor Neurons Drives C. elegans Forward Locomotion. Neuron 76, 750–761 (2012).

19. Hernandez-Nunez, L. et al. Reverse-correlation analysis of navigation dynamics in Drosophila larva using optogenetics. eLife 4, e06225 (2015).

20. Donnelly, J. L. et al. Monoaminergic Orchestration of Motor Programs in a Complex C. elegans Behavior. PLOS Biol. 11, (2013).

21. Silver, D. et al. Mastering the game of Go with deep neural networks and tree search. Nature 529, 484–489 (2016).

22. Silver, D. et al. Mastering the game of Go without human knowledge. Nature 550, 354–359 (2017).

23. Schrittwieser, J., et al. Mastering Atari, Go, chess and shogi by planning with a learned model. Nature 588, 604–609 (2020).

24. Mnih, V. et al. Human-level control through deep reinforcement learning. Nature 518, 529– 533 (2015).

25. Vinyals, O. et al. Grandmaster level in StarCraft II using multi-agent reinforcement learning. Nature 575, 350–354 (2019).

26. OpenAI et al. Dota 2 with Large Scale Deep Reinforcement Learning. http://arxiv.org/abs/1912.06680(2019) doi:10.48550/arXiv.1912.06680.

27. Wurman, P. R. et al. Outracing champion Gran Turismo drivers with deep reinforcement learning. Nature 602, 223–228 (2022).

28. Degrave, J. et al. Magnetic control of tokamak plasmas through deep reinforcement learning. Nature 602, 414–419 (2022).

29. Ibarz, J. et al. How to train your robot with deep reinforcement learning: lessons we have learned. Int. J. Robot. Res. 40, 698–721 (2021).

30. Haydari, A. & Yılmaz, Y. Deep Reinforcement Learning for Intelligent Transportation Systems: A Survey. IEEE Trans. Intell. Transp. Syst. 23, 11–32 (2022).

31. Haarnoja, T., et al. Soft actor-critic algorithms and applications. ArXiv Prepr. ArXiv181205905 (2018).

32. Christodoulou, P. Soft actor-critic for discrete action settings. ArXiv Prepr. ArXi*v191007207* (2019).

33. Wong, C.-C., Chien, S.-Y., Feng, H.-M. & Aoyama, H. Motion Planning for Dual-Arm Robot Based on Soft Actor-Critic. IEEE Access 9, 26871–26885 (2021).

34. Sarma, G. P. et al. OpenWorm: overview and recent advances in integrative biological simulation of Caenorhabditis elegans. Philos. Trans. R. Soc. B Biol. Sci. 373, 20170382 (2018).

35. Shorten, C. & Khoshgoftaar, T. M. A survey on Image Data Augmentation for Deep Learning. J. Big Data 6, 60 (2019).

36. Nikishin, E., et al. Improving Stability in Deep Reinforcement Learning with Weight Averaging. (2018).

37. Reinforcement Learning Resources — Stable Baselines 2.10.2 documentation. https://stable-baselines.readthedocs.io/en/master/guide/rl.html.

38. Bhardwaj, A., Thapliyal, S., Dahiya, Y. & Babu, K. FLP-18 Functions through the G-Protein-Coupled Receptors NPR-1 and NPR-4 to Modulate Reversal Length in Caenorhabditis elegans. J. Neurosci. 38, 4641–4654 (2018).

39. Riddle, D. L., Blumenthal, T., Meyer, B. J. & Priess, J. R. Mechanosensory Control of Locomotion. C. elegans II. 2nd edition (Cold Spring Harbor Laboratory Press, 1997).

40. Brandt, R., Gergou, A., Wacker, I., Fath, T. & Hutter, H. A Caenorhabditis elegans model of tau hyperphosphorylation: Induction of developmental defects by transgenic overexpression of Alzheimer’s disease-like modified tau. Neurobiol. Aging 30, 22–33 (2009).

41. Jospin, M. et al. A Neuronal Acetylcholine Receptor Regulates the Balance of Muscle Excitation and Inhibition in Caenorhabditis elegans. PLoS Biol. 7, e1000265 (2009).

42. Hollenstein, J., Auddy, S., Saveriano, M., Renaudo, E. & Piater, J. Action Noise in Off-Policy Deep Reinforcement Learning: Impact on Exploration and Performance. Trans. Mach. Learn. Res. (2022).

43. Andersen, R. A., Aflalo, T., Bashford, L., Bjånes, D. & Kellis, S. Exploring Cognition with Brain–Machine Interfaces. Annu. Rev. Psychol. 73, 131–158 (2022).

44. Tankus, A., Fried, I. & Shoham, S. Cognitive-motor brain–machine interfaces. J. Physiol. Paris 108, 38–44 (2014).

45. Sussillo, D., Stavisky, S. D., Kao, J. C., Ryu, S. I. & Shenoy, K. V. Making brain–machine interfaces robust to future neural variability. Nat. Commun. 7, 1–13 (2016).

46. Dong, X. et al. Toward a living soft microrobot through optogenetic locomotion control of Caenorhabditis elegans. *Sci*. Robot. 6, (2021).

47. Lee, J. B. et al. A Compressed Sensing Framework for Efficient Dissection of Neural Circuits. Nat. Methods 16, 126–133 (2019).

48. Kim, K. & Li, C. Expression and regulation of an FMRFamide-related neuropeptide gene family in Caenorhabditis elegans. J. Comp. Neurol. 475, 540–550 (2004).

49. Tandon, P. pytorch-soft-actor-critic. https://github.com/pranz24/pytorch-soft-actor-critic (2022).

50. alirezakazemipour/Discrete-SAC-PyTorch: PyTorch implementation of discrete version of Soft Actor-Critic. https://github.com/alirezakazemipour/Discrete-SAC-PyTorch.

