## Supplementary Information for "Improving animal behaviors through a neural interface with deep reinforcement learning"

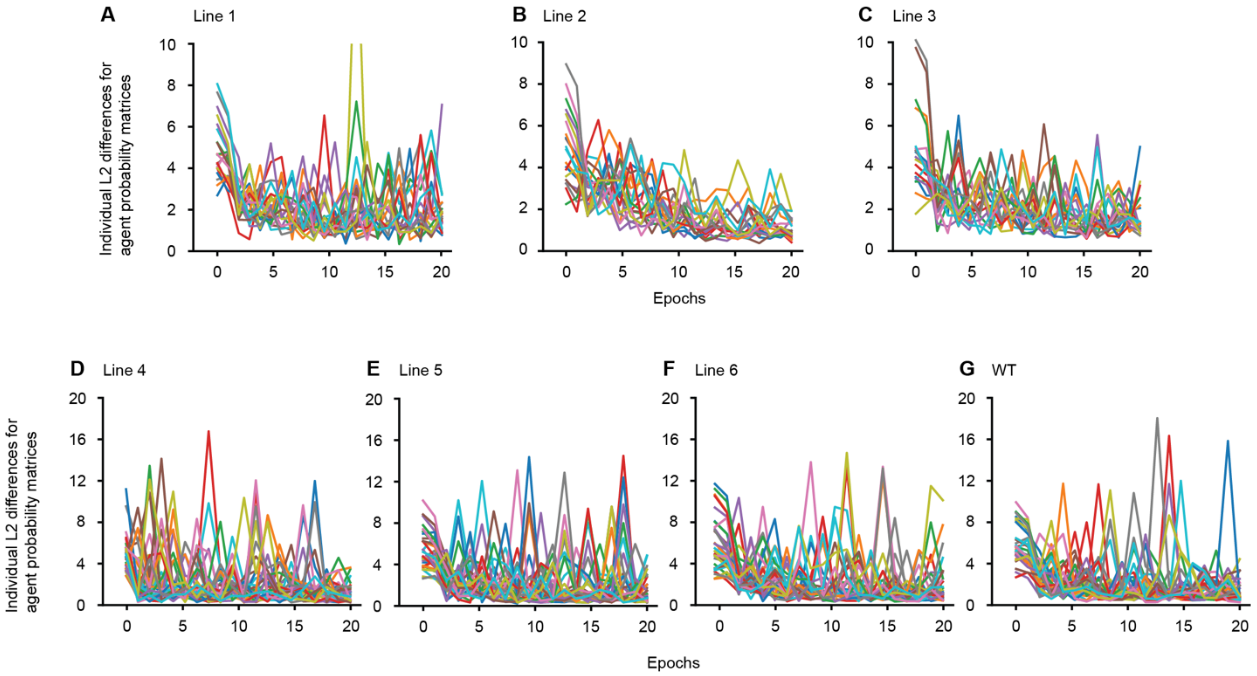


**Figure S1. Ensemble training stabilized performance.**

Individual agents during training were compared to ensembles after 20 epochs of training using a makeshift scoring metric. The metric was defined as the L2 difference between action probability matrices of the tested agent and the fully trained ensemble for each genetic line (as in Figure 2L-M).

(A-G) The individual errors for each agent in the ensembles for each transgenic line and wild type animals (G). Each color denotes a different agent. Note the relatively large variation among agents and instabilities, reflected by occasional sharp increases in error.

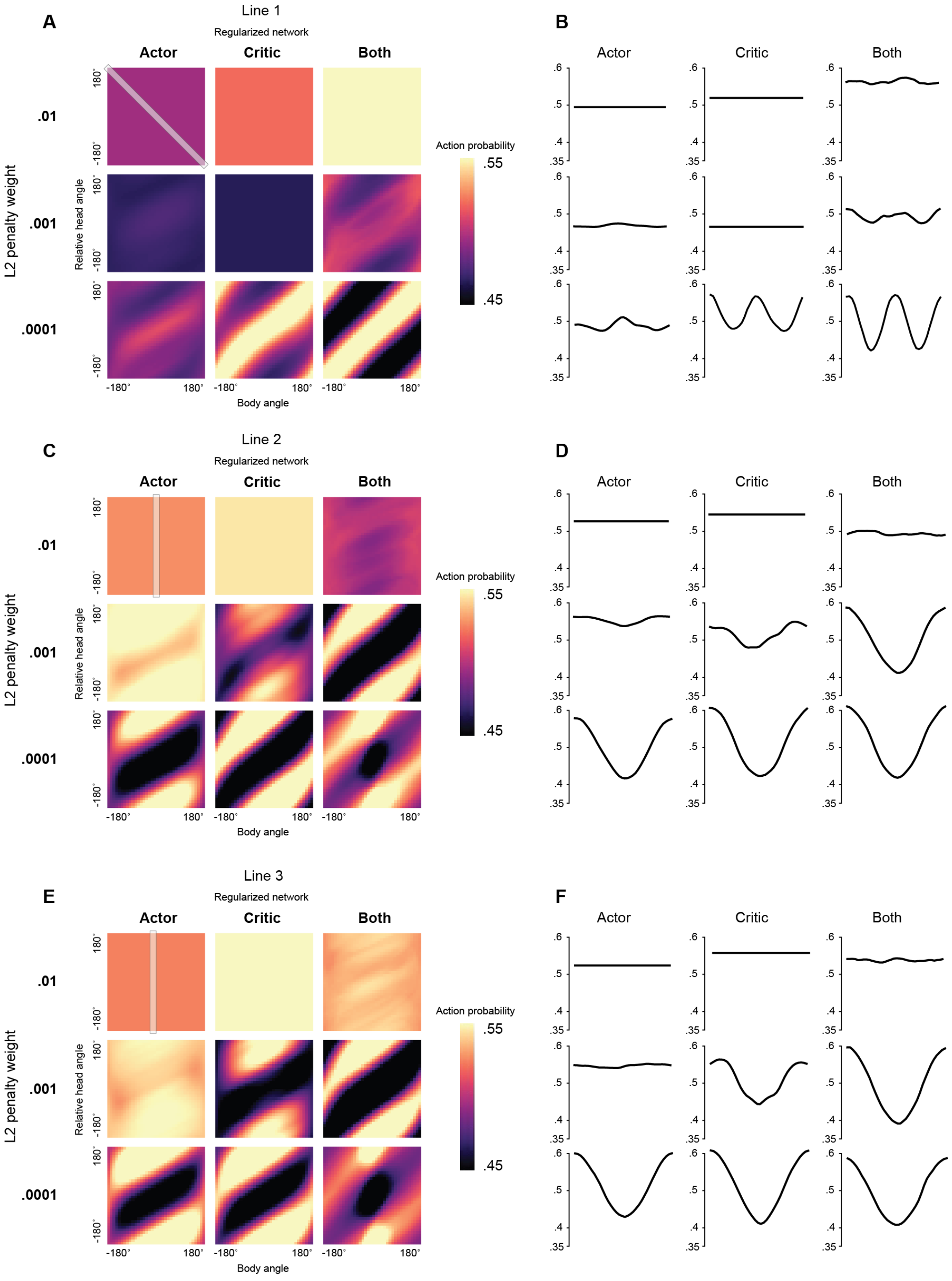


Figure S2. **L2 regularization did not noticeably improve learning and often made it worse.** (A, C, E) Ensemble action probability matrices (Figure 2J, 4D) using the same training methods as in agents used for evaluation (Methods), but for L2 weight penalties of 0.01, 001, 0.0001 over the whole network that the regularizer was applied to. These penalties are proportional to squared weight magnitudes.

(B, D, F) Traces of cross-sections highlighted in the top left matrix of (A, C, E). Traces follow cross-sections top-to-bottom along the x-axis.

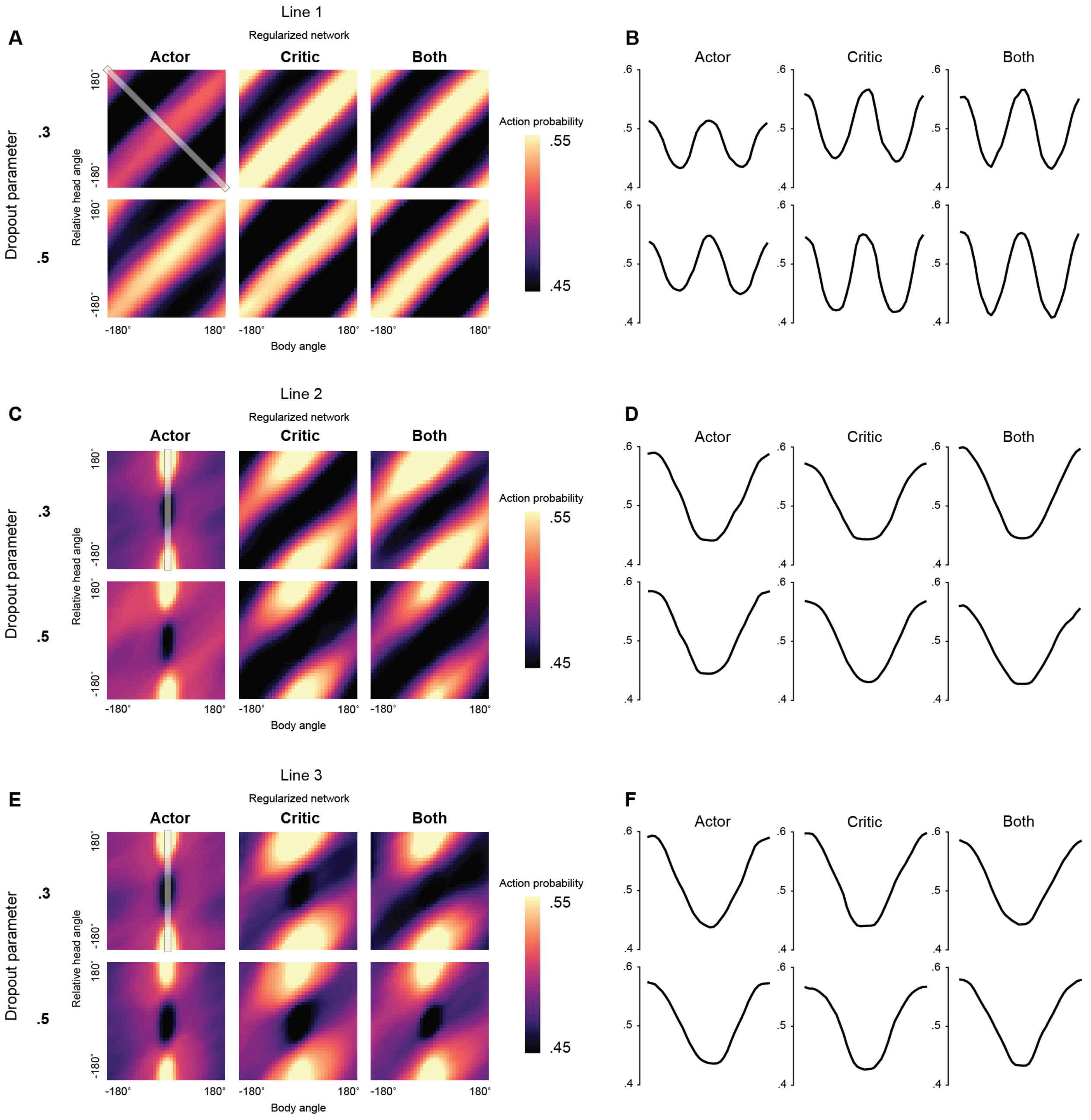


**Figure S3. Dropout did not noticeably improve performance when applied to actor networks, critic networks, or both.**

(A, C, E) Ensemble action probability matrices (following Figures 2J, 4D) using the same training methods as in agents used for evaluation (Methods), but for dropout rates of 0.3 or 0.5 over the whole network that dropout was applied to. These are rates at which neurons are randomly selected to not be part of the feedforward pass.

(B, D, F) show cross-sections highlighted in the top left matrix of (A, C, E). The x-axis for all plots follows the cross-sections top-to-bottom.

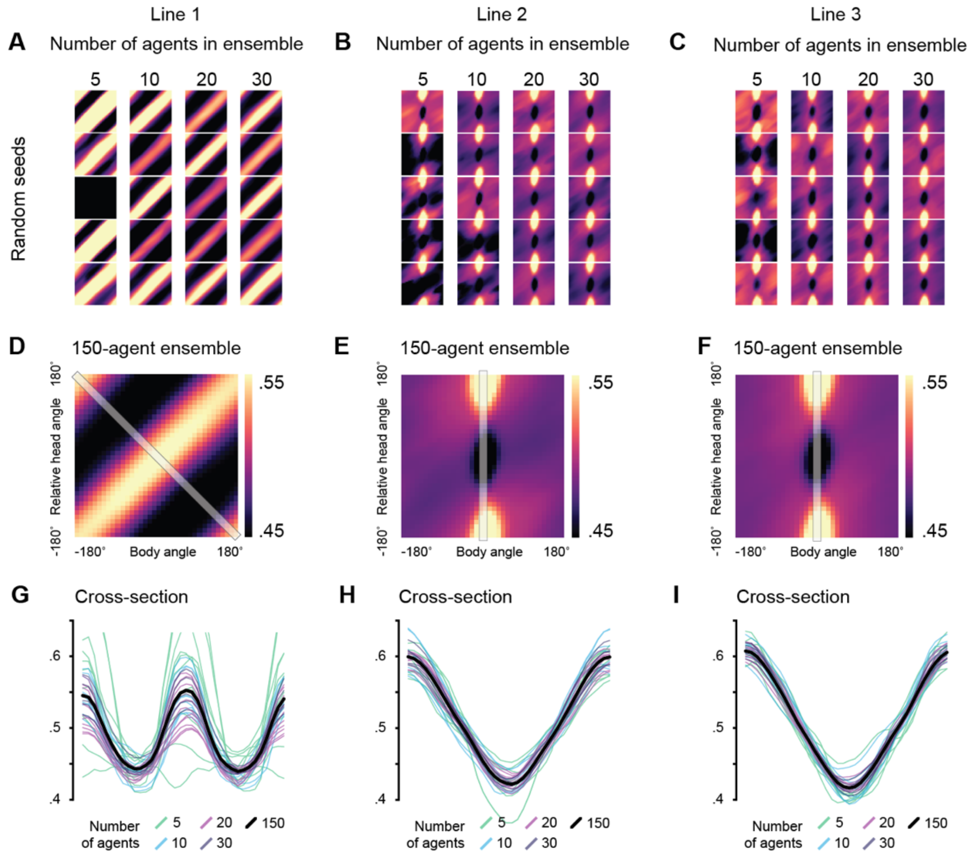


**Figure S4. Higher numbers of agents in an ensemble led to more consistent learned action probability matrices.** 150 agents were trained on the dataset for each genetic line.

(A–C) 5, 10, and 20 agents were drawn randomly without replacement from the 150-agent pool and their action probabilities averaged to form an ensemble. Data are for Lines 1-3. For the 30-agent column, the 150-agent pool was split into 5 equal parts to form ensembles.

(D–F) The entire pool of 150 agents was averaged to form one ensemble.

(G–I) Cross-sections of ensembles to show variability, taken from the highlighted strip in (D–F).

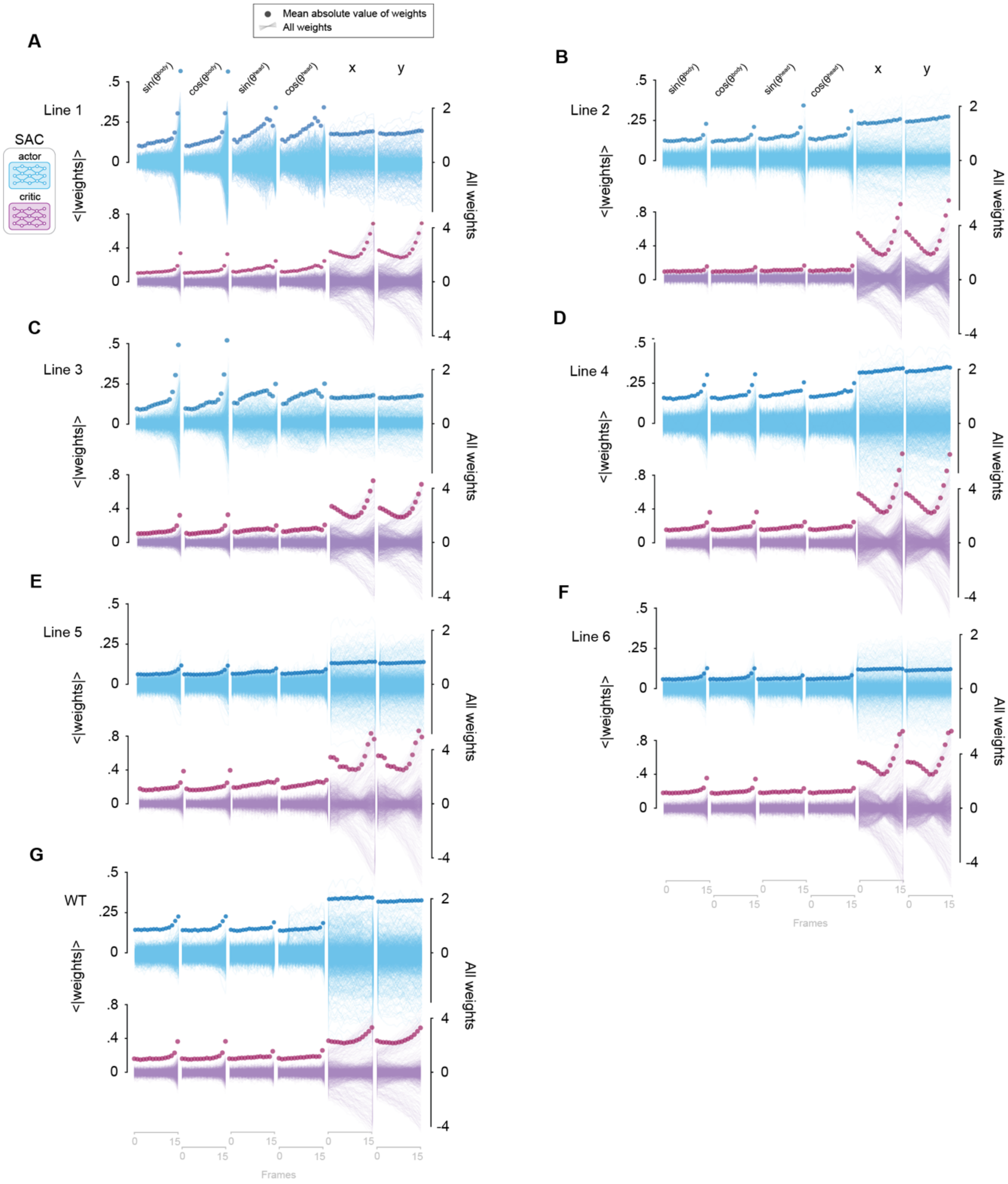


**Figure S5. Final timesteps for angle variables have larger average magnitudes for agents trained on all lines.**

(A-G) Weights in the first layer of the actor (top) and critic (bottom) networks in all 20 SAC agents of the ensemble trained on Lines 1-6 and wild type animals (G). Plotted as in Figure 2H. Raw weights are shown in lighter traces while the mean absolute values are plotted in darker circles to show weight trends.

| Agent parameters | Value |
| --- | --- |
| Temporal discount factor | 0.99 |
| Learning rate | 0.001 |
| Neurons per hidden layer | 64 |
| Layers per network | 2 |
| **Critic-only parameters** | **Value** |
| Target smoothing coefficient | 0.005 |
| Target update interview | 500 |

**Table S1. Agent parameters.**
Adam was used as an optimizer for actor and critic networks as well as automatic temperature tuning.

**Video S1. Line 1 with trained agent.** The video is sped up by 8x and shows that the animal is led to the target by the trained RL agent when coupled to neurons in Line 1. The target is a red circle and light flashes are denoted by a blue frame in this and all subsequent supplementary movies.

**Video S2. Line 1 with random light flashing.** The random frequency was matched to the average proportion of light-on measured across all standard evaluations in Figure 2G, calculated to be 0.4647. The video is sped up by 8x. When random flashes of light activate the neurons in Line 1, the animal is unable to reach the target.

**Video S3. Line 2 with trained agent.** The video is sped up by 8x and shows that after training, unlike in the control (Video S4), the RL agent excited neurons it was coupled to in Line 2 to direct the animal to the target.

**Video S4. Line 2 with random light flashing.** The random frequency was matched to the average proportion of light-on measured across all standard evaluations for Line 2 in Figure 3B, calculated to be 0.2896. The video is sped up by 8x. When random light flashes are used to activate neurons in Line 2 (see Figure 3A and Table 1 for line details), the animal is unable to reach the target.

**Video S5. Line 3 with trained agent.** The video is sped up by 8x and shows that the trained RL agent is able to learn appropriate patterns of light flashes based on the animal's posture to lead it to the target.

**Video S6. Line 3 with random light flashing.** The random frequency was matched to the average proportion of light-on measured across all standard evaluations for Line 3 in Figure 3B, calculated to be 0.3844. The video is sped up by 8x and again, with random flashes of light inhibiting neurons in Line 3, the animal does not reach the target.
